# A network for computing value homeostasis in the human medial prefrontal cortex

**DOI:** 10.1101/278531

**Authors:** Keno Juechems, Jan Balaguer, Santiago Herce Castañón, María Ruz, Jill X. O’Reilly, Christopher Summerfield

## Abstract

Humans and other animals make decisions in order to satisfy their goals. However, it remains unknown how neural circuits compute which of multiple possible goals should be pursued (e.g. when balancing hunger and thirst) and combine these signals with estimates of available reward alternatives. Here, humans undergoing functional magnetic resonance imaging (fMRI) accumulated two distinct assets over a sequence of trials. Financial outcomes depended on the minimum cumulate of either asset, creating a need to maintain “value homeostasis” by redressing any imbalance among the assets. BOLD signals in the dorsal anterior cingulate cortex (dACC) tracked the level of homeostatic imbalance among goals, whereas the ventromedial prefrontal cortex (vmPFC) signalled the level of homeostatic redress incurred by a choice, rather than the overall amount received. These results suggest that a network of medial frontal brain regions compute a value signal that maintains homeostatic balance among internal goals.

---

Canonical models in psychology, economics, and machine learning assume that decisions are made in order to maximise expected reward. One popular view proposes that estimates of reward for distinct stimuli or attributes (e.g. price and quality) are collapsed into a common value signal that motivates reward-based decisions^1^–^3^. In humans, BOLD signals^4^–^6^ in the medial orbitofrontal cortex (OFC) and ventromedial prefrontal cortex code for food items^7^,^8^, money^9^,^10^ and social stimuli^11^, suggesting the existence of a domain-general value code in this region. In non-human primates and rodents, neuronal firing rates^12^–^14^ in lateral and central orbitofrontal cortex (OFC) encode value signals for primary reinforcers. Moreover, humans and primates with medial OFC lesions fail to integrate multiple attributes of a stimulus when making decisions, implying that the OFC plays an active role in constructing composite value estimates for available stimuli^15^–^18^.

However, in natural environments, survival depends on the minimum level of any one of a number of competing internal needs. For example, a hungry animal needs food more than water, whereas the reverse is true for a thirsty animal. As the world changes dynamically, distinct internal assets (e.g. satiety or hydration) are continuously being depleted and replenished, so that homeostasis among internal resource levels needs to be monitored and maintained in neural circuits for valuation and choice^19^–^21^. The OFC is likely to play a role in evaluating stimuli in the context of internal goal states (e.g. satisfy hunger or quench thirst), because animals with OFC lesions continue to choose actions that lead to devalued outcomes, as if they were failing to maintain or follow currently active goals^18^,^22^. However, a key open question is how the brain evaluates which of multiple possible goals should be pursued at any one time, and how it dynamically updates goal values according to time-varying resource levels.

This question remains unaddressed because most previous studies have used paradigms (e.g. bandit or food choice tasks) that involve maximisation of a single asset, rather than those that require the balancing of multiple assets. Here, we built on a recent finding that medial OFC tracks the internal state given by cumulative satiety or wealth over a sequence of trials^23^,^24^. We reasoned that by encoding these signals, the OFC and interconnected areas of the medial prefrontal cortex might allow animals to compute ongoing levels of competing assets, and dynamically adjust choices to both replenish immediate needs and guard against future scarcity. We tested this using a task in which rewards depended on the minimum value of one of two assets. We report evidence for a new frame of reference for human neural value signals, with dACC encoding the imbalance among assets, and the vmPFC coding how this imbalance is redressed by a choice.

## Results

Participants (n = 21) performed a “virtual zookeeper” task in the fMRI scanner. The task involved managing a zoo that housed two assets (lions and elephants). On each trial, participants were offered the opportunity to expand their zoo by a variable number of animals (e.g. 4 lions or 2 elephants). Subsequently, trialwise reward was offered in proportion to the minimum number of animals in the zoo (e.g. if they had a total of 12 elephants and 8 lions in the zoo, they received 8 points) creating a pressure to redress any imbalance in the assets. To mimic the autocorrelation in natural environments, we created “streaks” in which numerically more of one species (4-6) were offered than the other (1-3); these numbers reversed with probability 0.3 on each trial. This manipulation ensured that the optimal policy was neither to always choose the more plentiful animal, nor to always satisfy immediate needs, but to vary these strategies according to the quality of the offer, the need to redress, and the time remaining in the block (see **Fig. 1a**).

**Figure 1.**
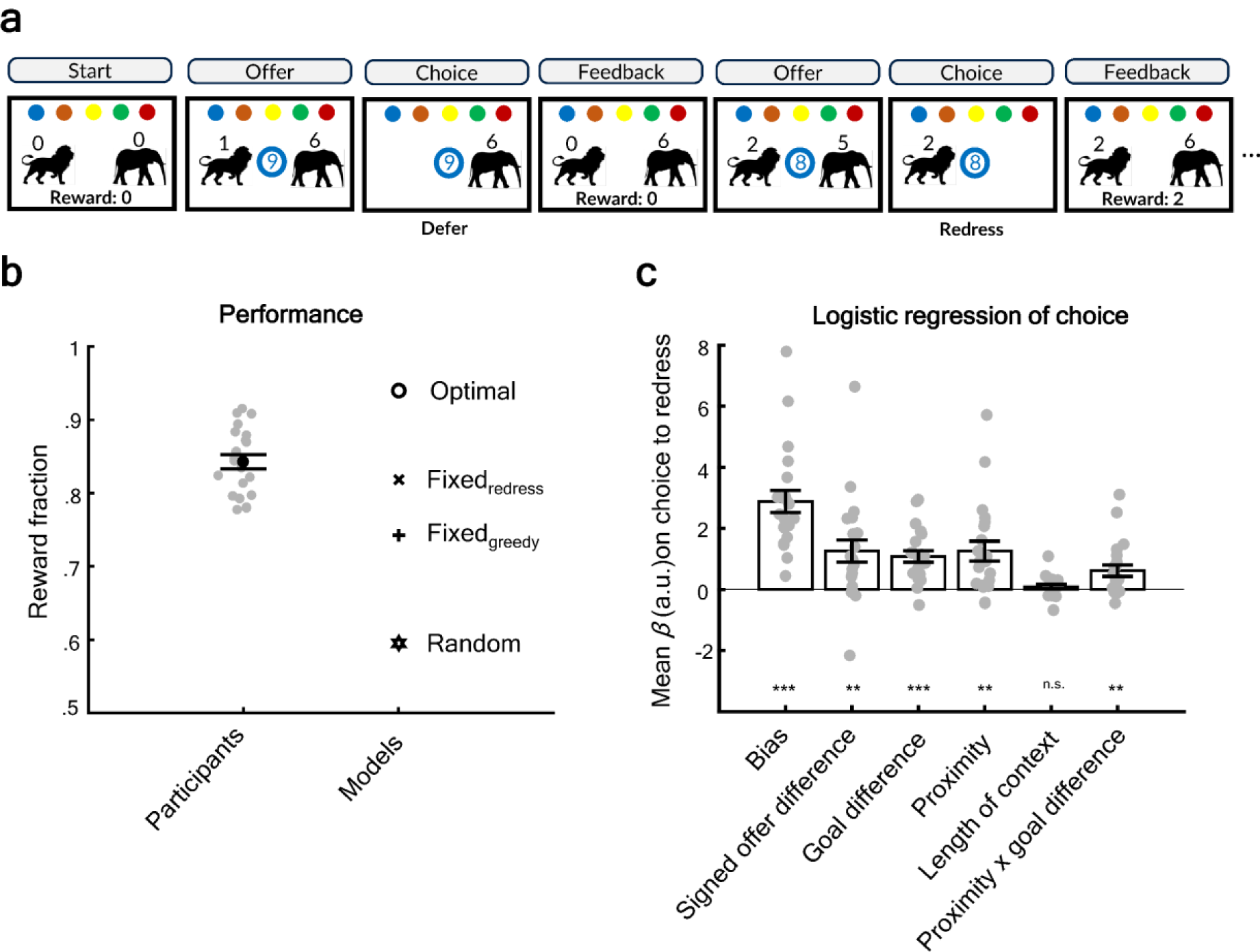
Task and behavior. (A) Task with 2 example trials from the first block in a scanner run. The number of animals on offer (“offer” screen), the number of animals chosen (“choice” screen) and the number of animals in the zoo (“feedback”) are shown numerically above a picture of a lion and elephant. Coloured circles (top) indicate the zoos (blocks) for the current run, with the central circle coloured according to the current block. The central number indicates the number of trials (including the current trial) remaining before the end of the block, i.e. the “countdown”. Trialwise reward was indicated on the feedback screen as the minimum of the two assets. Background screens were grey during testing, but are shown in white here for illustration. (B) Black circle indicates average earnings as fraction of maximum possible; black error bars are standard error of the mean (SEM) and grey circles show the data from individual participants. Other markers indicate averaged reward fractions for various models. (C) Beta coefficients from a logistic regression of various predictors on choice to redress. Bar height indicates mean beta, error bars are SEM, grey circles represent individual participant data. Significance was assessed using a two-tailed t-test against zero. *** is p < 0.001, ** is p < 0.01, * is p < 0.05.

### Behavioral data

Participants received 84.3% of the maximum attainable cumulative reward as calculated by numerical simulation (see Methods; **Fig. 1b**). This was significantly better than chance (mean: 59.6%, Wilcoxon sign-rank test, p < 0.001), than a fixed strategy of always choosing the higher offer (“greedy”; mean: 74.3%, p < 0.001), and than a model that used a fixed strategy of always satisfying immediate needs, assuming perfect memory for the tallies of animals in the zoo (“redress”; mean: 81.8%, sign-rank test, p < 0.03).

### Human choices are driven by both offer values and internal state

We used logistic regression to ask whether human choices were driven by both offers and needs (internal state). Choices were classed as either “redress” (mean = 80.85%, SD = 10.78%) when participants chose to acquire more animals of the species with the lower current tally and “defer” otherwise. Participants were strongly biased to choose to redress (t_20_ = 7.74, p < 1 × 10^−6^). However, this tendency was stronger when the *signed offer difference* was higher, i.e. when more of the currently minimal asset was offered; t_20_ = 3.36, p < 0.005), and also stronger when there was greater need to redress, i.e. when the level of imbalance among the two assets was higher (t_20_= 5.61, p < 1 × 10^−4^). We call this imbalance *goal difference*, and equate this with “internal” state or needs, because it was not shown to participants at the time of choice, but had to be maintained in memory over trials. Goal difference was defined as high asset minus low asset, which means that it was unsigned and its magnitude signals the pressure to redress. Together, these results show that participants were neither greedily choosing the highest offer, nor slavishly satisfying immediate needs, but rather trading off the relative offer values and their internal needs or goals when making decisions.

### The dACC encodes homeostatic imbalance

Behavioural analyses indicated that decisions to redress depended on the offer difference and the goal difference. Hence, in our first neural analysis (GLM1) we asked whether these quantities are encoded in blood-oxygenation-level dependent (BOLD) signals at the time of choice, i.e. time-locked to the offer. As previously described, clusters within the dorsomedial prefrontal cortex (dmPFC), extending into dACC (Brodmann area 32) correlated positively with the offer difference (see **Fig. 2b**; dACC peak: −6, 28, 38 (x,y,z), global peak: −14, 28, 50, cluster-false-discovery-rate-q (FDRq) < 1 × 10^−11^), as did those in the and bilateral insula (left peak: −34, 20, −2; right peak: 46, 20, 2; cluster – FDRqs < 0.02), bilateral inferior parietal lobule (iPL) and angular gyri (left peak: −46, −56, 30; right peak: 50, −56, 30, cluster – FDRqs < 0.005). However, the frame of reference of this neural signal was “unsigned” with respect to needs – it encoded the absolute difference in offer (no effects were observed for signed offer difference), **Table S1**.

**Figure 2.**
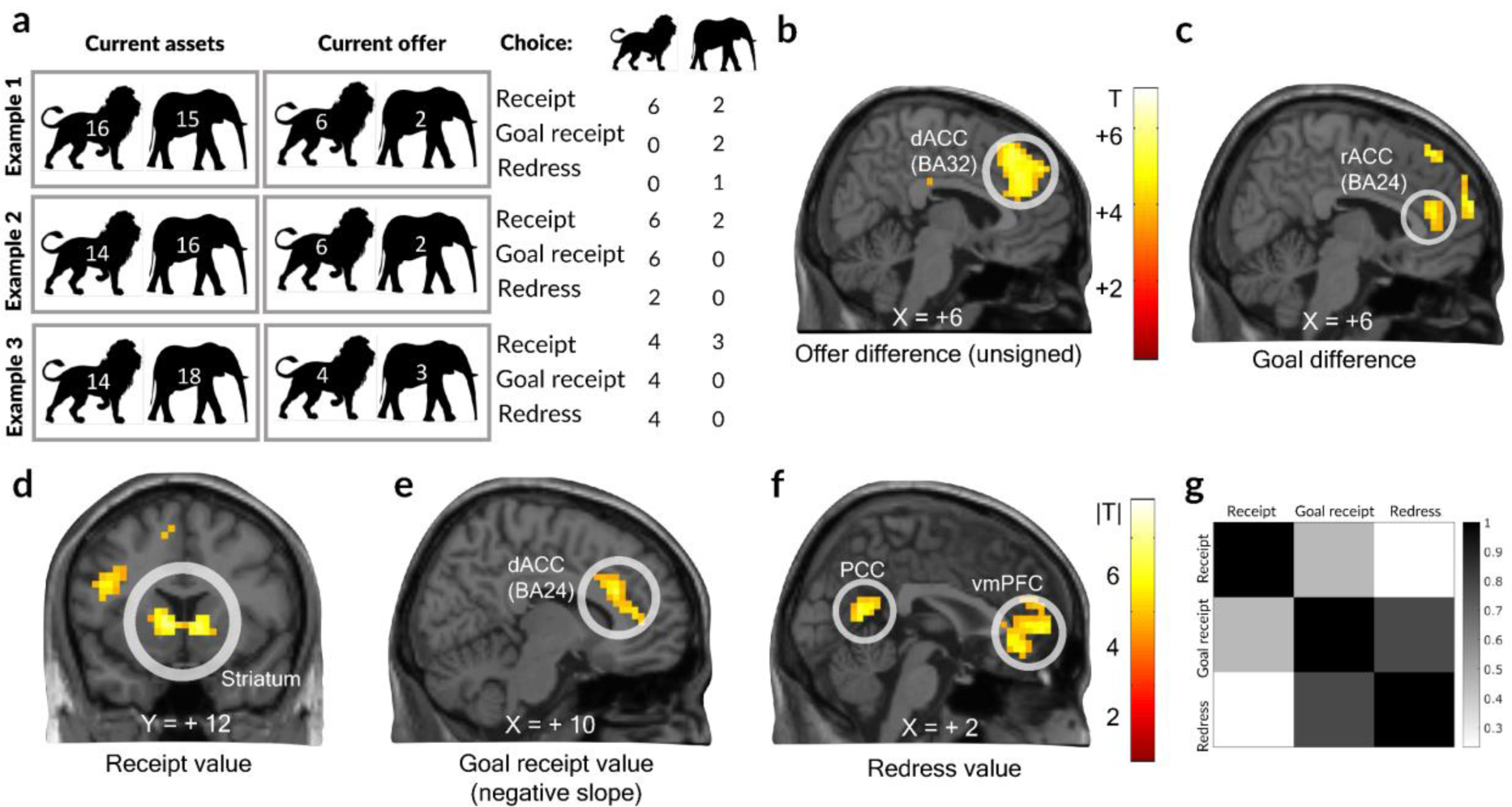
Representation of choice-relevant task features and value codes. (A) Illustration of the different value codes described in the paper for three example trials. The “current assets” panel depicts the state of assets before participants made a choice, whereas the “current offer” depicts the available choices. The panel to the right shows the values of the three value codes used for analyses depending on choice: Receipt value reflected the increase in the overall number of animals achieved by choice. Goal receipt value codes the receipt value in the frame of reference of the goal. Finally, redress value encodes the realized redress between the goals or, equivalently, the increase in trial reward. It is computed as min (receipt value, goal difference) if participants chose according to their needs, or zero otherwise. (B) Positive correlation between offer difference (high minus low) and BOLD signals in the medial PFC, sagittal slice at x = 6. (C) Positive correlation of goal difference with voxels on the same slice as in A). (D) Encoding of receipt value in voxels rendered on a coronal slice at y = 12. (E) Encoding of goal receipt value on a sagittal slice at x = 10 (F) Encoding of redress value on a sagittal slice at x = 2. Voxels were thresholded at p < 0.001, uncorrected. (G) Correlation matrix for the three value codes.

Critically, however, at the time of choice a cluster in the rostral ACC (rACC) positively encoded the internally maintained imbalance in assets (“goal difference”; BA 24 peak: 10, 40, 14, cluster – FDRq < 0.01; **Fig. 2c and Table S2**), together with a cluster in the ventral occipital lobe (peak: −34, −48, −18, cluster FDRq < 0.005). Neither offer difference nor goal difference was strongly correlated with reaction time (RT; r = 0.046, r = 0.006, respectively), and we found the same neural effects when controlling for RT in GLM1 (**Fig. S1**), so this finding does not reflect the confounding influence of time-on-task, but rather that the rACC tracks the level of imbalance among internal needs. Adjacent rostromedial regions of the PFC displayed a similar response to that observed in the rACC (**Fig. 2c**).

### VmPFC encodes a value redress signal

Outcomes were deterministic in our task, and so participants knew the consequences of their choices as soon as they were made. Previous research has reliably demonstrated that vmPFC encodes reward outcome, but left open the question of how current needs may impact this encoding. Thus, we designed our task to dissociate three possible frames of reference for the outcome signal that might be computed on each trial: the total number of animals received independent of need (receipt value), the number of animals received in the frame of reference of need (goal receipt value), and the level of redress (redress value). To illustrate, consider a participant with 10 lions and 12 elephants in their zoo who receives 3 more lions. The receipt value (and goal receipt value) would be 3, whereas the redress value would be 2. However, if the participant chose 2 more elephants, the receipt value would be 2, and the goal receipt value and redress value would be zero (see **Fig. 2a** for an illustration). By design, receipt value and redress value were only weakly correlated (r = 0.23, R^2^ < 0.06; correlation matrix **Fig. 2g**) and thus dissociable. Note that redress value does not equal the monetary reward received on a given trial, but rather encodes the expected *increase* in reward conditional on the choice.

We let these three value regressors compete for variance in the BOLD signal at the time of choice (GLM2). Receipt value positively activated clusters in the bilateral dorsal striatum (caudate nucleus), extending into ventral striatum (peak: −6, 12, 2; cluster – FDRq < 1 × 10^−6^; **Fig. 2d**). Goal receipt value was negatively correlated with an area in the dorsal ACC (BA 24; peak: 14, 28, 26; cluster – FDRq < 0.05 **Fig. 2e**). By contrast, redress value activated a cluster in the vmPFC (peak: 6, 52, 2, cluster – FDRq < 0.001; **Fig. 2f**) and the posterior cingulate cortex (peak: −6, −60, 14, cluster-FDRq < 0.005), two regions that are often found to coactivate in value-based decision tasks^4^ (see **Tables S3-5** for full results). This association between redress value and vmPFC signals was also observed when we coded redress value relative to unchosen redress, in line with the previous observation that the vmPFC codes for the value of a chosen relative to an unchosen alternative^25^,^26^. Moreover, when we took the conservative approach of first orthogonalising this relative redress value with respect to the other two predictors, ensuring that any effect of redress value must be over and above the other two value codes, we observed the same triple dissociation in dorsal striatum, dACC and vmPFC (**Fig. S2, Tables S6-8**). Finally, we used a non-circular ROI approach to test whether the vmPFC simply coded for the monetary reward observed on each trial by testing whether it encoded the level of the lower (reward-determining) asset at the time of choice, but found that this was not the case (t_20_ = 0.02, p > 0.98; GLM3). We thus conclude that vmPFC encodes the level of homeostatic redress incurred by a choice.

Together, thus, we find that the dorsal striatum, dACC and vmPFC fulfilled dissociable roles in value encoding in our task. The dorsal striatum encoded the number of animals received irrespective of needs (receipt value), whereas one region in the dACC (BA 24) participated in encoding goal difference (the need to redress assets) and the goal receipt value (the outcome value in the frame of reference of needs). In contrast, the vmPFC encoded the redress in homeostatic imbalance incurred by a given choice, i.e. the extent to which internal needs were returned to a balanced state by a given choice.

### Computational model and optimal policy

To ask how an agent should behave in order to maximise reward in our task, we used dynamic programming (DP) to compute the policy that will maximise expected future return over each zoo. DP searches through possible future states and chooses a reward-maximising action according to the expected future return, assuming optimal subsequent choices. Because it accounts for possible future offers up to the horizon defined by the end of the zoo, the DP algorithm will sometimes choose the immediately less needed option in order to maximize overall return. For instance, where an offer of the asset not immediately required is generous, it may be sensible to nevertheless choose that asset, to guard against future scarcity when the offer probabilities reverse. This policy will be less useful at the end of the block, because such future benefits are curtailed when the zoo ends.

Armed with this computational tool, we first computed the upper bound on reward shown in **Fig. 1b**, as reported above. Next, we asked whether humans, like the model, considered potential time horizon when choosing to defer an immediate redress in asset imbalance. We found that they did: as for the DP model, proximity to the end of a zoo (defined as 1/countdown) positively predicted redress responses (t_20_ = 3.74, p < 0.005) and that this tendency was stronger when the goal difference was greater (t_20_ = 3.13, p < 0.01). However, this was not driven by the overall length of the zoo alone, which failed to predict redress probability (t_20_ = 1.26, p > 0.22). Next, we evaluated choices made by the DP model using the same logistic regression approach. Despite some similarities in the betas obtained between humans and the DP model, the latter exhibited a stronger impact on *p(redress)* of both offer value and proximity to the end of the zoo (**Fig. S3)**.

These findings might be explained if humans adopt a strategy that is approximately optimal but myopic i.e. has insufficient search depth when computing expected future return. We thus reimplemented the DP model, but allowed the planning horizon (in steps) to vary as a free parameter, as well as the subjective reversal probability, and policy terms that allow for bias and choice variability. When we fit the output of this “myopic” DP models to participants choices using maximum likelihood estimation with five-fold cross-validation (one fold per scanner run), we found that, on average, participants estimated the reversal probability to be approximately 0.44 (SD = 0.33) and planned only 7.5 trials ahead (SD = 6.04), significantly lower than the theoretical value of 20. The overall model log-likelihood was −2047.2, which was lower than both the logistic regression model with proximity (7 parameters; LL = −2111.1) and without proximity (4 parameters; LL = −2189.2), and the DP model was strongly favoured by Bayesian model selection (exceedance probability = 0.99).

This best-fitting (but myopic) DP model was also able to reproduce several qualitative features of the data. First, it captured how the average probability of redress varied with proximity to the end of the block (**Fig. 3a**). Second, it captured how the average homeostatic need to redress (i.e. tolerance for disparity among the two assets in the zoo) varied as a function of time for blocks of different length, **Fig. 3b**. By contrast, a fully optimal DP model is more tolerant of larger goal difference values in the middle portion of the block, indicative of a longer time horizon for planning.

**Figure 3.**
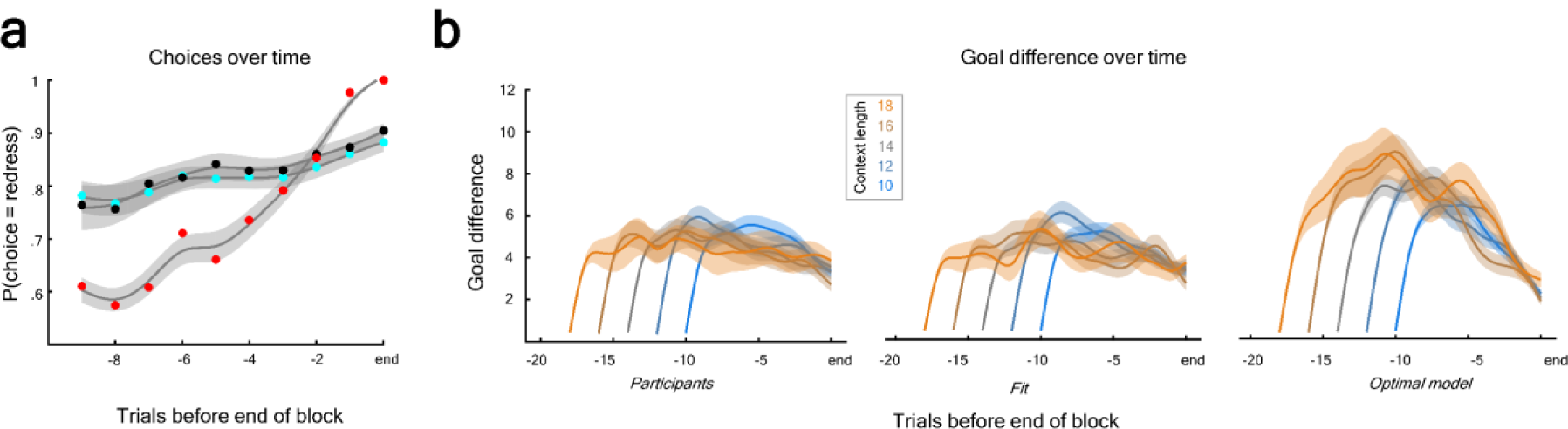
Comparison of optimal policy model and participants’ behaviour. (A) Plot of redress choices against countdown toward the end of a block for optimal DP model (red dots), human participants (black dots), and best fitting “myopic” DP model (turquoise dots). Circles are mean and shaded areas are SEM around the spline-smoothed means (for display purposes only). (B) Plot of goal difference against time for blocks (zoos) of different length. The three subpanels show data averaged across participants (left) the simulated fit of the myopic model (middle), and the optimal model (right panel). Solid lines are mean across participants/models, shaded areas are SEM. Curves are spline-smoothed for clarity of viewing.

In conclusion, the DP analyses revealed that although humans planned for future needs, they were suboptimal in two ways: firstly, they did not plan ahead sufficiently, and secondly, they over-estimated the reversal probability.

### Neural coding of time-varying pressure to redress

Returning our attention to the neuroimaging data, we asked whether BOLD signals covaried with the interactions of the two asset tallies and proximity, i.e. whether cortical regions reflected a time-varying pressure to redress. In GLM3 we found that the rACC (BA 24) region covarying with goal difference (i.e. the quantity that participants need to keep in homeostatic balance) also encoded the interaction of the higher asset and proximity (negatively; t_20_ = −2.16, p < 0.05), but not of the lower asset and proximity (t_20_ = 1.33, p > 0.19). Thus, encoding of the higher asset, and by extension goal difference, decreased over time in this region (**Fig. 4b)**. We also observed that a more posterior region in the dmPFC (GLM1; peak: −36, 16, 50, cluster – FDRq < 0.001), as well as bilateral angular gyri (left peak: −50, −48, 34; right peak: 54, −44, 38; cluster – FDRqs < 0.001), and dorsolateral PFC (left peak: −34, 20, 46; right peak: 30, 4, 62; cluster – FDRqs < 0.001, **Table S9**) encoded the main effect of proximity to the block end, independent of asset difference, as if they were tracking the available time remaining to redress (**Fig. 4a**).

**Figure 4.**
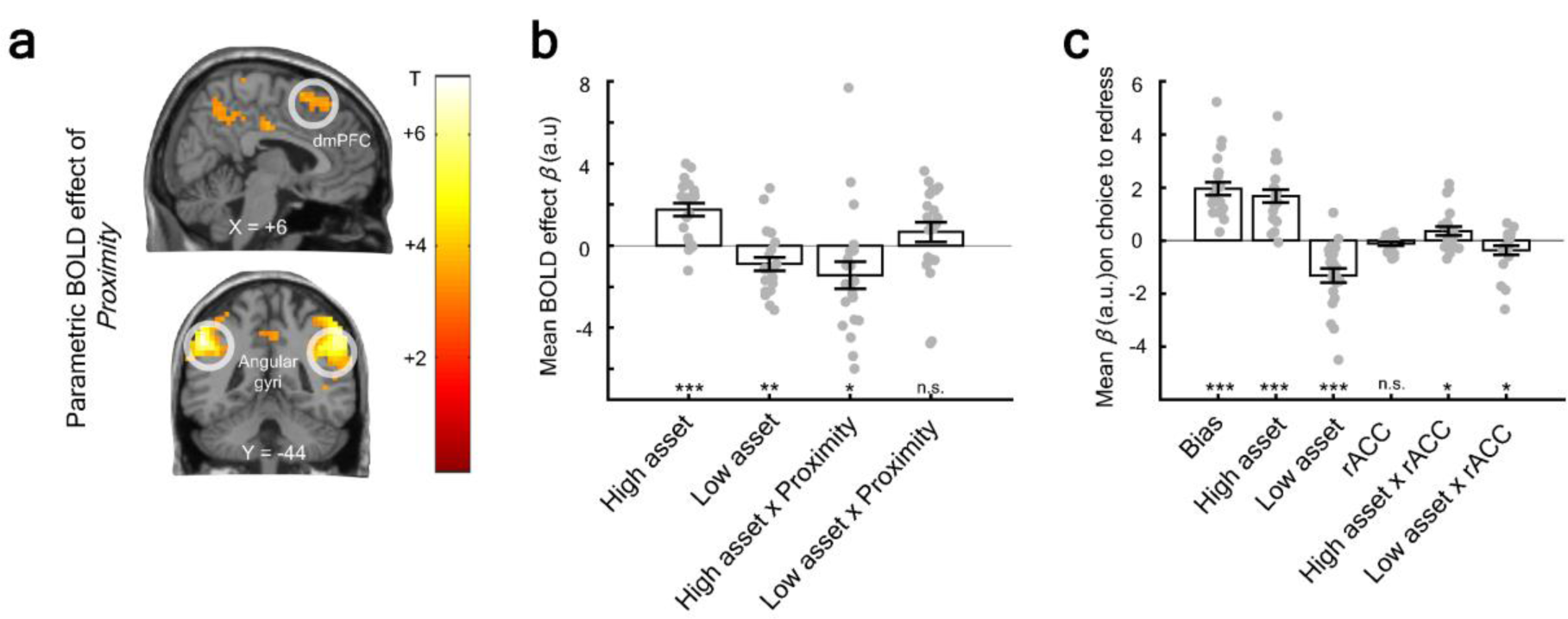
Neural effects of trial sequence. (A) Main effect of proximity on voxels rendered on a sagittal slice at x = 6 and a coronal slice at y = −44 from GLM1. Voxels were thresholded at p < 0.001, uncorrected. (B) Higher and lower asset encoding in leave-one-out ROI in rACC (BA 24; ROI defined from goal difference contrast in Fig. 2). Bar height indicates mean neural beta, error bars are SEM, grey circles represent individual betas. (C) rACC activity predicted choice to redress on a trial-by-trial basis. Bars are mean parameter estimates from logistic regression, error bars are SEM. Significance was assessed using a two-tailed t-test against zero. *** is p < 0.001, ** is p < 0.01, * is p < 0.05.

Finally, we asked whether single-trial activity (from GLM4) in the same rACC ROI (BA 24) had an impact on behaviour. We constructed a logistic regression model in which choices to redress were predicted by the two goal tallies, the rACC activity, and the interactions of rACC with the goal tallies. Choices to redress were negatively predicted by the lower tally (t_20_ = − 4.75, p < 0.0002), and this effect was accentuated when rACC signals were stronger (t_20_ = − 2.11, p < 0.05), whereas choices to redress were predicted positively by the higher tally (t_20_ = 6.69, p < 1 × 10^−5^), and this effect was also stronger when rACC was more active (t_20_ = 2.09, p < 0.05; **Fig. 4c)**. No main effect of rACC activity was observed (t_20_ = −1.72, p > 0.10). Thus, high rACC activity guided choices based on the current need to redress.

## Discussion

We asked humans to perform a value-guided decision task required them to balance levels of two possible assets (“value homeostasis”). We found that they did so strategically, basing their choices about whether or not to redress or defer on the value of current offers, knowledge of homeostatic imbalance, and the time horizon available for future redress. In other words, humans make choices that actively arbitrate among current goals, and plan for possible future scarcity. Our major contribution here is that considering this frame of reference for value signals sheds new light on the function of key brain areas involved in reward-guided decision-making.

Previous studies have identified human BOLD responses that code for the economic value of offers and outcomes in various frames of reference, observing for example that the vmPFC codes positively for the value of a chosen option and negatively for an unchosen option, with the reverse pattern observed in the dACC^4^. Other, less well-established frames of reference for value signals have been observed, for example that vmPFC^27^,^28^ and dACC^29^ encodes value relative to an average or default stimulus, or that they respectively signal the value of a proximal choice, and the need to switch to a new context of known average value^30^–^32^. However, these studies make a critical assumption that is uncharacteristic of the real world: that choices are made to maximise momentary return given a single, monolithic value function, rather than arbitrating among multiple possible goals which depend on internal needs or cumulative assets. By contrast, here we start with the prediction that animals evolved not simply to maximise momentary outcomes but to maintain a homeostatic balance among multiple internal needs. This is the case not just for primary reinforcers that promote satiety and hydration, but also for more complex, abstract needs, as when a busy academic balances the goals of conducting impactful research and providing effective teaching. Our work sheds new light on the function of three regions previously implicated in value-guided choice: the dorsal and parts of the ventral striatum, vmPFC and dACC. We report a clear triple dissociation between these regions, which is striking given that previous studies have largely suggested that they jointly compute values of stimuli, actions and goals^33^,^34^.

Firstly, the striatum responded to outcome magnitude (i.e. number of animals received), but unlike cortical regions, it did so in a fashion that was insensitive to internal needs – the signal went unmodulated by estimated cumulative assets. Previous studies have suggested that where a single asset (e.g. money) is to be maximised, both vmPFC and striatum seem to code the long-run average value of an action, and the value computed via tree search through future states or outcomes^4^,^33^,^35^. These common value signals in cortical and subcortical regions may reflect the fact that animals use a mixture of model-free and model-based information when making decisions^33^,^34^,^36^. However, our work instead suggests that cortical systems compute values in the frame of reference of ongoing internal needs, whereas the striatum codes for overall receipt independent of internal needs.

Unlike previous studies, we found that the vmPFC did not code for outcomes *per se*, either in the frame of reference of choices or contexts. Rather, it codes for homeostatic redress – that is, the extent to which any given choice restores the imbalance among internal needs. This signal remained the best explanation of vmPFC signals even when we partialled out the actual outcome received in number of animals, or the trialwise monetary reward that motivated participants to perform the task. We note that in our task, as in a natural environment where wellbeing is determined by the minimum of multiple possible assets, redress value is inevitably identical with the change in reward or hedonic value. This presumably explains the ubiquitous observation that vmPFC correlates positively with reward outcome when a single asset is being optimised, and why this signal is often observed to be modulated by the local context provided by average value^28^,^37^,^38^.

These findings build on our previous work implicating the human vmPFC in coding an internal, unsignalled representation of cumulative assets in a way that maximises a specific goal (maintaining net positive aggregate reward), over and above any momentary hedonic value that arises from choices^23^. However, the current work additionally implies *why* the brain keeps track of cumulative assets – because this allows any imbalance among internal needs to be redressed by future choices. This helps understand a perplexing paradox in the decision literature – how is it that neural signals in the vmPFC code ubiquitously for rewards, but lesions to the vmPFC incur only subtle deficits where decisions involve maximisation of a single asset^39^? The vmPFC outcome signals ubiquitously observed in fMRI studies may reflect the tracking of redress value to one of multiple potential goals – obtaining food, or money, or maximising accuracy – rather than computing the relative strength of one offer over another. Indeed, in studies of navigation the vmPFC signal also ramps up over multiple steps that are made towards a destination, presumably because each step redresses the distance between current and goal state^40^,^41^. Thus, we would expect patients with vmPFC lesions to arbitrate effectively between offers of differing value but to fail to allocate decisions strategically over the long term, precisely the pattern that is observed in both humans and monkeys^15^,^39^.

Perhaps the most interesting neural effects we report, however, are two dissociable value signals in adjacent, and partially overlapping, ACC regions, which collectively imply that this region participates in choosing which goal to pursue at any one time. Firstly, a cluster in BA24 responded to the current imbalance among assets, i.e. the extent to which one asset needed to be replenished in order to match another. Furthermore, the amplitude of this signal was predictive of choices to redress. Previous studies of reward-guided decisions have emphasised that the dACC codes positively for the relative value of an “unchosen” option– i.e. that which was foregone when a choice was made^26^,^42^ – or that it signals the need to switch away from a current context towards a new, potentially richer, source of rewards^30^,^43^. Our suggestion that the rACC and adjacent dACC encode homeostatic imbalance jointly explains these findings, which respectively occur during instances of heightened value imbalance between one stimulus and another, or one context and another. Moreover, lesions of the dACC lead monkeys to switch away from a currently rewarding task, as if they had difficulty maintaining goal value^44^,^45^. Secondly, an adjacent region in a more dorsal part of BA 24 signals the extent to which an asset that is needed is replenished by a choice (goal receipt value). We note that this signal is required to compute the vmPFC redress signal, when combined with knowledge of cumulative assets. Together, these findings argue for a critical role for the ACC in computing which of multiple possible goals should be pursued at any one time.

The choice of whether to satisfy an immediate need, or to build up less pressing resources to offset future scarcity, is a ubiquitous problem for humans and other animals^46^. Critically, this choice depends on the time horizon over which choices can be made: an optimal agent with knowledge of the time horizon will tolerate an asset imbalance early in the block, where it can be later redressed if opportunities change, but respond to pressure to redress later in the block. Human participants were less tolerant of imbalance than the optimal model, which can be explained if they are somewhat myopic in their planning. Note, however, that this is akin to, and indistinguishable from a parameter discounting future rewards in favour of short-term gain. Nevertheless, reversal probability, planning horizon and an alternative discount parameter would all predict that participants chose to redress more than was optimal, especially at the early stages of the block. Consistent with this view, the BA24 ROI that coded for goal difference did so in a fashion that varied with proximity to the end of the block. However, the direction of this effect was somewhat surprising to us, in that goal difference encoding decreases with time. It is possible that participants had a bias to redress towards the end irrespective of the size of the goal difference – thus, goal difference encoding may not be needed to drive choice towards the final few trials in a block. Nevertheless, this finding requires further corroboration in future studies.

Our findings may more generally have implications for understanding health and wellbeing in humans. We assume that wellbeing is related to the minimum among current needs, and find evidence that neural circuits for valuation and choice have evolved to maintain a balance among needs via goal-directed choice. Disorders of valuation and choice, such as depression, might be associated with failures of a system that attempts to maintain homeostatic imbalance, such as an exaggerated representation of goal difference, or a failure to update internal asset estimates when they are redressed.

## Online Methods

### Participant details

Twenty-two healthy volunteers with no history of psychiatric or neurological disorder participated in the study. One participant was excluded from analyses due to scanning equipment failure, leaving N = 21 (5 male, mean age = 23.6, SD = 2.77). All participants gave written informed consent and the study received approval from the ethics board of the University of Granada. Participants were compensated for their time with€25 plus a performance-based bonus (mean =€7.15, SD =€1.42, range:€4.52 to€11).

### Task details

Participants performed a task that involved managing a virtual zoo which housed two types of animals: lions and elephants. Each zoo began empty and animals were offered over a block lasting a variable number of trials (10-20), before a new zoo began. Participants encountered 5 zoos on each of 5 scanner runs, with each run lasting a total of 65 trials. On each trial, participants were first shown a fixation cross (150 ms), followed by a “choice” screen (2.5 seconds) in which they were offered a variable number of lions or elephants. This was indicated by an arabic number above a drawing of the animal on the left and right of the screen. There were always a high (4-6) offer of one animal and a low (1-3) offer of the other. The assignment of high or low offers to each animal was autocorrelated over trials, reversing with a probability of 0.3, which led to “streaks” in which one animal could be harvested in more abundance. Participants indicated their choice by pressing a button with their left or right index finger (for options on the left and right, respectively). The side on which each animal was offered varied randomly across trials. Once a choice was made, the unchosen option disappeared from the screen, leaving only the chosen option (“response” screen), and this lasted the remainder of the trial (2.5s minus reaction time), followed by a blank screen for a uniformly jittered interval (1.5-4.5 seconds). Subsequently, a feedback screen (1 second) indicated the number of animals of each species in the zoo (the “assets”) as a number above the respective drawing. Trial-wise reward was indicated below the drawings and was directly proportional to the lower asset (e.g. it was 5 if there were 10 elephants and 5 lions in the zoo). Inter-trial intervals were drawn uniformly between 2s and 6s (1-3 time of repetition; TR).

Throughout the run, five coloured discs just below the top of the screen were shown, representing in the 5 zoos ordered from left to right in time. Each zoo had a colour, and the colour of the current zoo was indicated by a central circle that appeared on the “choice” and “response” screens. Colours were drawn randomly on each run and were only included to aide participants in grouping trials into discrete blocks. During the “choice” and “response” phases of the trial, a central number was displayed within the circle, also in a colour matching the colour of the current zoo. This number counted down the trials still remaining in the current zoo (including the current trial). At the conclusion of a block, its cumulative earnings were displayed on a “bonus” screen and this number was shown underneath the corresponding coloured disc on all subsequent trials. A lottery at the end of each run displayed to participants which of these bonuses was chosen as extra payment. Each block had equal probability of being selected in the lottery.

#### Training and instruction

Participants completed one run immediately before scanning to familiarize themselves with the task. They were told that if lions were currently the high offer, they were more likely to be the high offer on the following trial, but that this was not certain and this contingency could reverse unpredictably. However, they were not told the exact reversal probability (0.3). They were also told that they would receive the sum of the lottery values across runs after the end of the experiment.

### Analysis of behavior

All analyses were carried out using MATLAB 8.6 (R2015b) and custom in-house code.

Choices were coded as “redress” when participants chose the offer corresponding to the currently lower asset in their zoo and “defer” if they chose the other option. All behavioural results discussed in the main text are based on this dichotomy (but see **SI** for behavioural analyses that classed choices as “greedy” with respect to the offers on the screen; **Fig. S4**). Proximity was coded as 1/countdown.

### Logistic regression

The dependent measure for the logistic regression was the binary variable 1=redress, 0=defer. We fit the models to individual subjects using z-scored regressors in five-fold cross-validation (one for each scanner run). We then averaged the betas across runs for each subject and used a two-tailed one-sample t-test to determine whether the group mean of each parameter differed significantly from zero. Trials where the goal difference was zero were excluded from analysis (n = 1071 out of N = 7150 trials, of which almost half were first trials in a new block).

The first logistic regression model contained the following regressors: a constant term, signed offer difference (need option minus not-needed option), absolute asset difference, and overall length of block. The second logistic regression model added proximity and the interaction of goal difference and proximity.

Model fits were compared using five-fold cross-fitting via Matlab’s fitglm function, summing log-likelihoods across participants and trials. In addition, we performed Bayesian model selection to compute the exceedance probabilities for each model, which measures whether a given model was more likely to be the true model than all competitor models^48^.

### Dynamic Programming model

We developed a Markov model to derive the optimal solution for the task. The model’s state space was given by the goal difference, the number of trials left in the block, the two offers in the frame of reference of the lower asset, and the reversal probability (constant for all models, except when fitting to participants). The model was optimized using the following dynamic programming formula

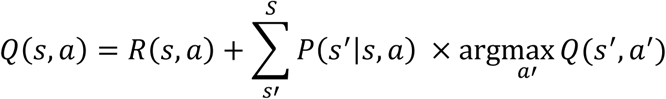

Where *s* and *s’* are the current and successor state, *a* and *a’* are the actions taken in each state, Q(s,a) is the value of taking action *a* in state *s*, and P(s’|s,a) is the probability of transitioning from state *s* to state *s’* by choosing action *a*, and R(s,a) is the reward obtained in the current state. Thus, the model chose the action which maximized the sum of immediate reward (R(s,a)) and the expected value of the best action in all subsequent states. As the task had a finite horizon, we do not include a discount factor. We fit the model with different reversal probabilities (from 0 in increments of 0.05 to 1, where 0.3 was the true value) in order to compare it to participants’ behaviour. As the model computed the expected value for both choosing to redress and choosing to defer, we computed the difference between the two and passed it through a sigmoid to allow for noisy decisions. The output of this sigmoid – p(redress) – was then fit to participant’s behaviour with a weighting parameter and intercept using maximum likelihood estimation. Maximum possible reward was calculated by permuting across all possible combinations of choices in each sequence and finding the highest reward score.

### FMRI Data Acquisition and pre-processing

MRI data were acquired on a 3T Siemens scanner. T1 weighted structural images were recorded directly prior to the task using an MPRAGE sequence: 1×1×1mm^3^ voxel resolution, 176×256×256 grid, TR = 1900ms, TE = 2.52ms, TI = 900ms. Each fMRI image contained 32 axial echo-planar images (EPI) acquired in descending sequence with a voxel resolution of 3.5 mm isotropic, slice spacing of 4.2mm, TR = 2000ms, flip angle = 80, and TE of 30ms. 1850 EPI images were recorded per participant, resulting in a scanning time of around 62 minutes. Data were pre-processed using SPM12 (Wellcome Trust, London). As EPI acquisition used a descending sequence, images were corrected for slice time acquisition with the middle slice (at TR/2 = 1 s) as reference to minimize interpolation errors^47^. Scans were realigned to the first scan within each session. The anatomical scan was co-registered to the mean functional image. Anatomical scans were normalized to the standard MNI152 template brain and smoothed with a 4mm full width half maximum Gaussian kernel. The functional EPI images were then normalized and smoothed with a full width half maximum Gaussian kernel of 8mm.

### FMRI data analysis

Data were analysed using SPM12 and custom scripts. Data from the five scanning sessions were concatenated and constants identifying each run were added manually to the GLM in order to account for potential differences in mean activation and scanner drift. Stimulus onsets were incremented by 1s to account for slice time correction to the middle slice, which occurred at TR/2 = 1s after stimulus onset^47^. Micro time onset was adjusted to the first slice. All results reported came from GLMs in which the canonical haemodynamic response function was convolved with a delta function (i.e. with zero seconds duration) time-locked to event onset (e.g. offer or feedback screen). The six vectors of motion parameters derived from pre-processing were included as nuisance regressors. Automatic orthogonalization was disabled. Group-level contrasts were constructed as simple t-contrasts using subject-level contrast images as input. Throughout this paper, we report only clusters that survived false-discovery-rate-correction (FDR) for multiple comparisons^49^,^50^ unless otherwise noted. This avoids the potentially inflated false positive rates at the cluster level^51^. Cluster extent was set to 10 or more voxels and voxel-wise threshold to p < 0.001, uncorrected. Data were visualized using the xjview toolbox for SPM (http://www.alivelearn.net/xjview).

Our analyses are based on 4 GLM models. All 4 GLMs included two predictors of interest (onset of the choice screen, onset of the feedback screen) and two nuisance predictors (trials on which the goal difference was zero, and all no-response trials). GLM1 additionally included the following parametric regressors time-locked to the offer (i.e. at the time of choice): congruency (whether the high offer was of the currently needed type), response hand (left vs. right), signed and unsigned offer difference, sum of the offers, proximity, and goal difference. For completeness, we also included the following parametric modulators on trials with zero goal difference: response hand, unsigned offer difference, offer sum, proximity. Finally, goal difference (post-choice) was included as parametric modulator time-locked to the feedback screen. GLM2 included parametric modulators for the receipt value, goal receipt value, and redress value, as well as proximity, all time-locked to the onset of the offer. For completeness, we also included chosen value and proximity on trials with zero goal difference, and goal difference and cumulative value at feedback. GLM3 included parametric modulators corresponding to the two zoo tallies (up to that trial) their interactions with proximity, and their increases, as well as proximity. These modulators were time-locked to the onset of the choice screen. These regressors were z-scored to allow comparisons between their beta values. For completeness, we also included chosen and foregone receipt value on trials with zero goal difference, as well as goal difference and total sum of assets at feedback. GLM4 modelled each trial and feedback screen as a single event without any parametric modulation.

#### Region of interest extraction

ROI were extracted using a leave-one-subject-out procedure to avoid potentially circular inference. A mask was created for each participant using the first-level contrast images from all other participants (i.e. leaving out the current participant). ROI were then constructed by taking all significant voxels (p < 0.001, uncorrected) within the relevant region (approximately given by xjview’s database, e.g. within medial PFC). We extracted one rACC ROI from the “goal difference” contrast from GLM1 and one vmPFC ROI from the “redress value” contrast from GLM2.

### Materials and Correspondence

Behavioral data and custom code to reproduce figures can be accessed on our GitHub profile at the point of publication. All requests for further information, code, or fMRI data should be addressed to Mr. Keno Juechems, Department of Experimental Psychology, University of Oxford

## Author contributions

KJ, CS, and JXO designed the study and interpreted the results, KJ, JB, SHC, MR, and CS collected the data, JB provided code for fMRI analyses, KJ analysed the data and provided code, KJ and CS wrote the paper.

## Acknowledgments

This work was supported by European Research Council to CS; and the Economic and Social Research Council studentship to KJ (ES/J500112/1); and a Wellcome Trust 4-year-PhD grant to SHC (0099741/Z/12/Z). We would like to thank Nathaniel Daw and his lab for useful suggestions, and all members of the Summerfield lab, especially Andreas Jarvstad, for helpful discussions from initial idea to publication, and Hannah Tickle, Hildward Vandormael, and Vickie Li for assistance collecting the data.

